# Differences In Neural Activity, But Not Behavior, Across Social Contexts In Guppies, *Poecilia Reticulata*

**DOI:** 10.1101/265736

**Authors:** Eva K. Fischer, Sarah E. Westrick, Lauren Hartsough, Kim L. Hoke

## Abstract

Animals are continually faced with the challenge of producing context-appropriate social behaviors. In many instances, animals produce unrelated behaviors across contexts. However, in some instances the same behaviors are produced across different social contexts, albeit in response to distinct stimuli and with distinct purposes. We took advantage of behavioral similarities across mating and aggression contexts in guppies, *Poecilia reticulata,* to understand how patterns of neural induction differ across social contexts when behaviors are nonetheless shared across contexts. While these is growing interest in understanding behavioral mechanisms in guppies, resources are sparse. As part of this study, we developed a neuroanatomical atlas of the guppy brain as a research community resource. Using this atlas, we found that neural activity in the preoptic area reflected social context, whereas individual differences in behavioral motivation paralleled activity in the posterior tuberculum and ventral telencephalon (teleost homologs of the ventral tegmental area and lateral septum, respectively). Our findings suggest independent coding of social salience versus behavioral motivation when behavioral repertoires are shared across social contexts.

**Summary statement:** Activity in distinct brain regions reflects behavioral context versus social motivation in a in which behavioral repertoires are shared across social contexts (Trinidadian guppies, *Poecilia reticulata*).

## Introduction

Producing behaviors appropriate to the current social context is a central challenge for animals, requiring the integration of both internal and external cues. Integrating cues from conspecifics is particularly critical, as interactions with potential mates, competitors, and offspring generally require distinct behavioral repertoires and physiological states. However, in some instances, similar behaviors are deployed across social contexts, albeit with distinct purposes. For example, frogs, birds, rodents, and humans produce similar vocalizations in the context of mate attraction and territory defense (Catchpole, 2008; Portfors, 2007; Wells, 2007). While behaviors are the same across contexts in these cases, they are produced in response to distinct stimuli, indicating that divergent sensory inputs are converted into similar behavioral outputs while presumably simultaneously maintaining information concerning social salience. Thus, the neural underpinnings promoting context-dependent behaviors are particularly intriguing in these situations, where seemingly identical behaviors are performed across clearly distinct social contexts.

Trinidadian guppies, *Poecilia reticulata,* perform similar, highly stereotyped behaviors during mating and aggressive interactions, and provide an excellent system in which to examine how social salience is reflected in the brain when behaviors are shared across contexts. Trinidadian guppies have become a model system in evolutionary ecology and behavior. Owing to extensive work in both the wild and the lab, much is known about the environmental cues influencing behaviors and the ultimate adaptive significance of these behaviors in this species (Houde, 1997; Magurran, 2005). Guppies are live-bearing fish with internal fertilization, and males spend the majority of their time in pursuit of females. Male guppies display a stereotyped courtship behavior known as a sigmoid display, during which they orient themselves perpendicular to a female, assume the characteristic S-shape that gives the display its name, and quiver their bodies. As an alternative to these overt courtship displays, male guppies also attempt to gain fertilizations by forced/sneaky copulation. In this case, males approach females from behind and below and thrust their gonopodium (intromittent organ) forward toward the female’s genital pore. In addition, males will bite, head-butt, and body-slam females to get their attention (reviewed in Houde, 1997). Despite obvious functional differences, male guppies perform a strikingly similar set of behaviors in aggressive competitions with other males: courtship displays, forced copulation attempts, and physical contacts serve to achieve successful copulations in a mating context and to establish dominance hierarchies in an aggressive context (Houde, 1988; Houde, 1997).

Given overlap at the behavioral level, how are male guppies nonetheless attuned to obvious contextual differences between mating and aggressive interactions? In the present study, we examine the neural mechanisms mediating behavior across social contexts in guppies. We begin by characterizing behavior in mating versus aggressive contexts. Using an atlas of the guppy brain we constructed, we next describe patterns of neural activation associated with mating versus aggressive contexts across 13 brain regions. We focus our analysis on brain regions mediating social behaviors that are evolutionarily ancient and functionally conserved across vertebrates (the Social Decision Making Network, SDMN; O’Connell and Hofmann 2011). Finally, we combine behavioral measures with neural activity data to understand associations between neural induction and behavioral output. The results of this study build resources for future work examining neural mechanisms of behavior in this increasingly popular system and provide insights into how distinct brain regions can contribute to social context versus social motivation.

## Materials and methods

### Animals

All fish used in this study were sexually mature males from a single lab-reared population derived from the Marianne River Drainage in the Northern Range Mountains of Trinidad. Fish were housed on a 12:12 hour light cycle (lights on 7:00am to 7:00pm) and fed a measured food diet once daily. Fish received Tetramin™ tropical flake paste and hatched *Artemia* cysts on an alternating basis. All animal husbandry, experimental methods, and tissue collection proceedures were approved by the Colorado State University Animal Care and Use Committee (Approval #12-3818A).

### Behavior

Males were assigned to one of three experimental groups: aggression, mating, or isolation (n=10 per group). Fish were assayed concurrently in sets of three per day, with one representative from each experimental condition. Fish were placed in individual 2.5 liter tanks on the afternoon preceding behavioral trials. Behavioral trials were conducted the following morning, two hours after lights-on and lasted 60 minutes thereafter. In the aggression condition, two unfamiliar males were introduced into the focal male’s tank at the start of the trial. In the mating condition, two unfamiliar females were introduced. In the isolation condition males remained isolated in their tanks throughout the trial. Behaviors were continuously recorded by two independent observers using JWatcher™ software. Each observer watched either the aggression or the mating condition at alternating 15 minute intervals, such that behaviors recorded for each social condition were evenly distributed among observers. Tanks were isolated from one another by opaque barriers so fish could not see one another and behavioral trials were conducted behind a blind with tanks lit from above to reduce visibility of the observers to the fish.

We followed behavioral protocols previously established in our lab to define and record behaviors (Fischer et al., 2016). Previous work in our lab demonstrates that guppies perform similar behaviors in aggression and mating contexts and so the same behaviors were scored in both contexts. These included the number and duration of sigmoid displays, the number of forced copulation attempts, the number of gonopodial swings, the number of aggressive contacts (biting, head-butting, body slamming, tail slapping), and the number of posturing incidents (when fish line up nose to nose).

### Tissue collection

Whole brains were collected immediately following behavioral trials. Guppies were anesthetized by rapid cooling, followed by decapitation. Whole heads were fixed in 4% paraformaldehyde at 4°C overnight and then transferred to 30% sucrose for dehydration. Following dehydration, whole heads were embedded in mounting media (Tissue-Tek^®^ O.C.T. Compound, Electron Microscopy Sciences, Hatfield, PA, USA), rapidly frozen, and stored at -80°C until cryosectioning. Heads were sectioned in the coronal plane at 14µm, thaw mounted serially onto charged slides (Superfrost Plus, VWR, Randor, PA, USA), and stored at -20°C until immunohistochemical staining.

### Immunohistochemistry

We used a phosopho-S6 antibody that targets phosphorylated ribosomes (pS6; Life Technologies, Carlsbad, CA, USA) to assay neural activity. Ribosomes become phosphorylated following changes in electrical activity in neurons and the pS6 antibody therefore acts as a general marker of neural activation, akin to immediate early genes (Knight et al., 2012). As time course is critical for experiments involving immediate early genes and can vary across species, we assessed staining intensity at three timepoints (30, 60, and 90 minutes) in sample guppy tissue prior to the experiment and chose the 60-minute time point based on these preliminary results (data not shown).

We followed standard immunohistochemical procedures for antibody staining to label pS6positive neurons. Briefly, we quenched endogenous peroxidases using a 3% H_2_O_2_ solution, blocked slides in 5% normal goat serum diluted in 1X phosphate-buffered saline (PBS) and 0.03% tween for one hour, and then incubated slides in the anti-pS6 primary antibody (Life Technologies, Waltham, MA, USA) at a concentration of 1:500 in blocking solution overnight at 4°C. The following day, we incubated slides in secondary antibody (Jackson ImmunoResearch, West Grove, PA, USA) at a concentration of 1:200 in blocking solution for two hours, incubated slides in an avidin-biotin complex (ABC) solution (Vector Laboratories, Burlingame, CA, USA) for two hours, and treated slides with 3,3'-diaminobenzidine (DAB; Vector Laboratories, Burlingame, CA, USA) for five minutes to produce a color reaction. Slides were rinsed in 1X PBS prior to and following all the above steps. Finally, slides were rinsed in water, dehydrated in increasing concentrations of ethanol (50%, 75%, 95%, 100%, 100%), and coverslipped with Permount (Fisher Scientific, Hampton, NH, USA).

### Microscopy and cell counting

To reliably quantify neural activity across candidate brain regions, we created a guppy brain atlas (Supplemental Materials). We examined coronal brain sections of multiple male and female guppies stained using cresyl violet to assess morphology. We identified brain regions using neuroanatomical information from other fishes (Anken and Rahmann, 1994; Munchrath and Hofmann, 2010; Wullimann et al., 1996) and have made the atlas freely available online.

We photographed brain tissue at 20x magnification on a light microscope (Zeiss AxioZoom, Zeiss, Oberkochen, Germany) attached to a camera (ORCA-ER, Hamamatsu, San Jose, CA, USA) and analyzed cell counts from photographs using FIJI software (Schindelin et al., 2012). We outlined and measured focal brain regions (Fig. 1) and counted all stained cells within a given region. All regions extended across multiple sections and we quantified cell number for each region in all possible sections. We counted cells in only a single hemisphere per section.

**Figure 1.**
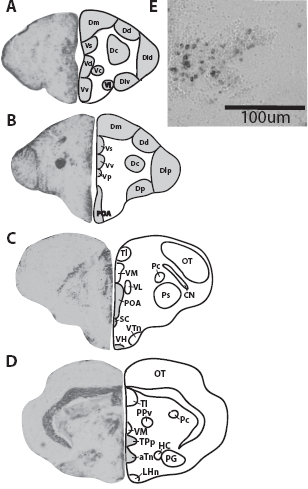
Overview of neuroanatomical atlas used for quantification of neural activation. **(A – D)** Representative sections of the telencephalon. We quantified neural induction in 14 candidate regions (shaded grey). **E)** Representative image of pS6 immunohistochemical staining from Vv. aTn = the anterior tuberal nucleus; Dc = central part of the dorsal telencephalon; Dd = dorsal part of the dorsal telencephalon; Dld = dorsolateral part of the dorsal telencephalon; Dlv = ventolateral part of the dorsal telencephalon (presumptive homologue of the mammalian hippocampus); Dm = medial part of the dorsal telencephalon; Dp = posterior part of the dorsal telencephalon; POA = preoptic area; TPp = the posterior tuberculum; Vc/Vd = dorsal and central parts of the ventral telencephalon (presumptive homolog of the mammalian nucleus accumbens and striatum); VH = ventral hypothalamus; VI = lateral part of the ventral telencephalon (presumptive homologue of the mammalian lateral septum); Vv = ventral part of the ventral telencephalon (presumptive homologue of the mammalian lateral septum).

### Statistical Analysis

We tested for the influence of social context (aggression versus mating) on behavior using generalized linear mixed models with a negative binomial distribution appropriate for count data with unequal variances. We tested for differences in the number of times each behavior was performed during the 60-minute trial. In addition, we summed the counts of all behaviors into a single total behavioral metric to assess overall behavioral activity. We chose this approach as (1) summing preserves the count nature of the original data, and (2) we have previously shown – and confirmed with exploratory analyses here – that the correlations between behaviors are not consistent across contexts and therefore the same principal components or factors cannot accurately summarize behavioral variation across experimental groups.

We used linear mixed models with a negative binomial distribution to test for differences in neural activation based on social context (aggression versus mating versus isolation). The model included social context and brain region as independent variables and the number of pS6-positive cells in each section as the dependent variable. We included fish identity as a random effect to control for repeated sampling within and among regions. As we expected that only some regions would show activation differences based on social context, we used Tukey-corrected posthoc tests to examine differences in a region-specific manner.

Finally, we tested whether activation in some regions predicted individual differences in behavior. To do this, we ran separate models for each brain region, in which the number of pS6positive cells per region, social context, and their interaction predicted the total number of behaviors during the trial. We excluded fish from the isolation treatment for this analysis because we had no behavioral data for these fish. We chose to use our total behavioral metric because (1) we wanted to use a metric that reflected general behavioral motivation, and (2) this approach increased the power of the statistical analysis. All statistical analyses were performed in SAS 9.4 (SAS Institute, Cary, NC, USA).

## Results

Fish performed similar behaviors between aggressive and mating contexts, and we observed no statistically significant differences (p<0.05) in the number of single or overall behaviors performed across mating and aggressive contexts (Table 1).

We assessed differences in neural activation among social contexts (aggression, mating, isolation) by quantifying region-specific pS6 immunoreactivity. Social context influenced neural induction of pS6 in a brain region-specific manner (context*region: F_24,1778_=3.70, p<0.0001). Posthoc analyses (Table 2) revealed that region-specific differences were significant in the preoptic area (POA; F_2,1778_=6.47, p=0.0016): Fish in the mating context had higher pS6 activation in the POA compared to fish in the aggression (t=2.79, adj p=0.015) or isolation (t=3.35, adj p=0.0024) context, which did not significantly differ in their extent of neural activation (t=0.53, adj p = 0.8569) (Fig. 2).

**Table 1.** Summary of tests for group differences in behavior.

| **behavior** | **df** | **F** | **p** |
| --- | --- | --- | --- |
| sigmoid display | 1,18 | 3.89 | 0.064 |
| sigmoid duration | 1,18 | 3.74 | 0.069 |
| gonopodial thrust | 1,18 | 3.60 | 0.074 |
| gonopodial swing | 1,18 | 4.39 | 0.051 |
| contacts | 1,18 | 0.96 | 0.334 |
| posturing | 1,18 | 4.25 | 0.054 |
| total behaviors | 1,18 | 1.80 | 0.197 |

**Table 2.** Posthoc analyses of regional differences in neural induction.

| brain region | df | F | p |
| --- | --- | --- | --- |
| aTn | 2,1838 | 1.57 | 0.208 |
| Dc | 2,1838 | 2.12 | 0.121 |
| Dd | 2,1838 | 0.37 | 0.688 |
| Dld | 2,1838 | 0.98 | 0.376 |
| Dlv | 2,1838 | 2.28 | 0.103 |
| Dm | 2,1838 | 0.23 | 0.793 |
| Dp | 2,1838 | 2.51 | 0.082 |
| POA | 2,1838 | 6.79 | **0.001** |
| TPp | 2,1838 | 1.67 | 0.188 |
| Vd | 2,1838 | 0.13 | 0.878 |
| VH | 2,1838 | 1.27 | 0.280 |
| VI | 2,1838 | 0.67 | 0.512 |
| Vv | 2,1838 | 0.49 | 0.616 |

**Figure 2.**
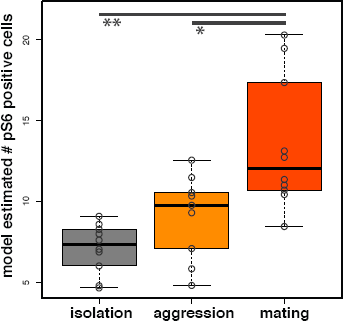
Differences in neural induction in the POA. Male guppies who experienced the mating context showed increased neural activation as compared to isolated fish and those in the aggression context. Model-estimated numbers of pS6 positive cells per section, rather than raw cell counts, are plotted. Circles overlaid on boxplots represent estimated, average cell number for each individual. Asterisks indicate p-values: * ≤ 0.05, ** ≤ 0.01.

Finally, we tested for both context-dependent and context-independent relationships between pS6 immunoreactivity and behavior. pS6 induction in the posterior tuberculum (TPp: F_1,15_=8.14, p=0.012) and the lateral part of the ventral telencephalon (Vl: F_1,14_=4.88, p=0.044) were positively associated with behavior in a context-independent manner (Fig. 3). We found no significant context-dependent relationships (Table 3).

**Table 3.**
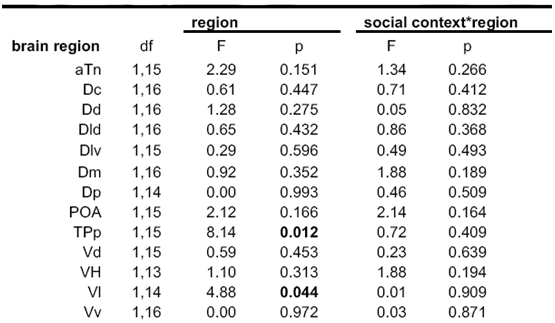
Test of brain region dependent and independent associations between neural induction and behavior.

**Figure 3.**
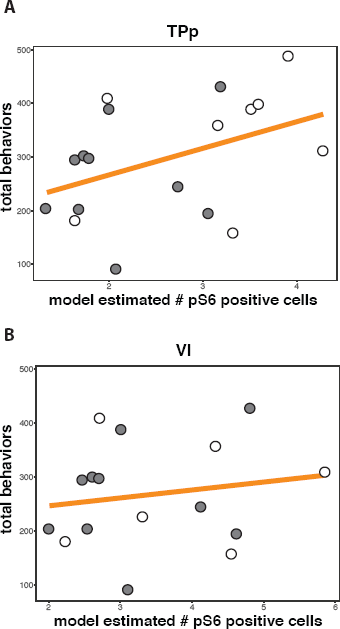
Relationship between neural activity and behavior. Increased neural induction in **(A)** the TPp and **(B)** Vl was associated with an increased number of total behaviors independent of social context (aggression = white circles, mating = grey circles). Model-estimated numbers of pS6 positive cells per section for each individual, rather than raw cell counts, are plotted.

## Discussion

In the present study, we sought to shed light on the neural mechanisms mediating social behavior across contexts and to understand how context is reflected at the neural level when behaviors are shared across distinct social contexts. To do so, we characterized neural activity patterns associated with shared behavioral repertoires across mating and aggression contexts in adult male guppies.

Adult male guppies performed highly similar behaviors in competitive and mating contexts. Extensive work has demonstrated that male guppies use alternative behavioral strategies under differing environmental conditions (e.g. predation threat, female receptivity, light levels, food availability) (reviewed in Houde, 1997; Magurran, 2005), yet competitive interactions between males have rarely been considered (but see Houde, 1988; Fischer et al., 2016). Our data suggest that, under the same environmental conditions, guppies direct similar behaviors with similar frequencies towards competitive (i.e. male) and mating (i.e. female) social partners (see also Fischer et al., 2016). Given previous work documenting behavioral flexibility in guppies in response to contextual factors including the presence of predators, abiotic differences, the reproductive state of females, and a choice between same sex or opposite sex social partners (reviewed in Houde, 1997; Magurran, 2005), we suggest that alternative contextual factors are more important in modifying the frequency and type of behaviors males direct at male versus female conspecifics.

Despite the lack of behavioral differences between contexts, mating and aggression present distinct challenges and opportunities for males and we therefore expected social context to be encoded at the neural level. Indeed, we observed that neural activation increased in the preoptic area (POA) of males following interactions with female – but not male – conspecifics. The identification of neural activity differences in the POA associated with distinct social contexts is in line with the functionally-conserved role of the POA in regulating social and sexual behavior in fish and other vertebrates.

The POA is an evolutionarily ancient brain region that is largely hodologically, molecularly, and functionally conserved across vertebrates. POA activity in general, and in particular arginine vasotocin (the teleost homolog of the mammalian nonapeptide arginine vasopressin; Foran and Bass, 1999; Semsar et al., 2001; Larson et al., 2006; Godwin and Thompson, 2012; O’Connell et al., 2012; Ramallo et al., 2012) and sex-steroid hormone (Forlano et al., 2005; Forlano et al., 2010; Munchrath and Hofmann, 2010) signaling play a central role in regulating social and sexual behaviors (reviewed in O’Connell and Hofmann 2011). In male fish, electrical stimulation of the POA has been show to increase reproductive behaviors (Demski and Knigge, 1971; Satou et al., 1984; Wong, 2000) and decrease aggression (Demski and Knigge, 1971), while ablation of the POA eliminates spawning reflexes (Macey et al., 1974). Although male guppies perform the same behaviors across mating and aggression contexts, copulation is only possible in interactions with females and differences in POA activation may be related to this critical distinction. Indeed, increased POA immediate early gene activity has been observed in response to sexual – but not aggressive – interactions in voles (Wang et al., 1997) and birds (Alger and Riters, 2006; Alger et al., 2009; Riters and Ball, 1999). Moreover, although no data are available outside of mammals, the POA has been shown to be necessary for ejaculation in rodents and monkeys (Malsbury, 1971; Slimp et al., 1978). As internal fertilization is rare in fish, guppies provide a unique system in which to more explicitly examine mechanistic convergence of ejaculation behavior in future. Our observations add to the body of work documenting widespread functional conservation of the POA in regulating social behavior across vertebrates and demonstrate a role for differential POA activity across social contexts, even when the behaviors performed across those contexts are shared.

In addition to identifying brain regions reflecting social context differences, we asked whether neural activation in some regions predicted behavior, either in a context-dependent or independent manner. We did not identify any regions with context-dependent associations, but did identify two brain regions in which levels of activation were positively associated with the number of behaviors performed in both aggressive and mating contexts: the posterior tuberculum (TPp) and the lateral part of the ventral telencephalon (Vl). Homologies of the TPp and Vl to mammalian brain regions remain somewhat contentious (Northcutt, 2008; Vargas et al., 2009). In fish, the TPp is generally proposed to be homologous to part of the mammalian ventral tegmental area (VTA), and Vl, together with the ventral part of the ventral telencephalon (Vv), is the putative homolog of the mammalian lateral septum (LS) (Rink and Wullimann, 2001; Rink and Wullimann, 2004). Acknowledging ongoing controversy concerning these homologies, we interpret our findings in view of what is currently known about these regions from fish and other vertebrates.

The VTA is a central component of the dopaminergic reward system, which plays a critical role in evaluating stimulus salience and motivating behavior. The LS receives projections from the VTA as well as the hippocampus, the hypothalamus, and the midbrain (Meibach and Siegel, 1977; Staiger and Nürnberger, 1989; Swanson and Cowan, 1979). It plays a role in both sexual and social behavior, in particular in the context of social memory, social recognition, and evaluating stimulus novelty (Bielsky et al., 2005; Dantzer et al., 1988; Landgraf et al., 1995; Liebsch et al., 1996; Maeda and Mogenson, 1981). In short, both regions are critical in evaluating, responding to, and retaining social information. Moreover, the connections between these two regions are thought to be particularly important in modulating goal-directed behavior (reviewed in O’Connell and Hofmann 2011). Given that all behaviors we assayed are typically directed at other individuals, and that we found group independent associations between neural activity and behavior in these particular regions, we propose that increased neural induction in these regions is associated with increased social behavioral motivation across contexts.

Though the majority of functional studies on behavioral motivation come from mammals, substantial work in song birds provides a particularly interesting comparison, as birds perform very similar singing behaviors across social contexts, analogous to the similarities in behavior across contexts we observed in guppies. Moreover, songbirds’ motivation to sing is clearly distinct from their ability to sing, as the latter is controlled by a well-defined set of song nuclei in the brain, while the former is controlled largely by the POA, the VTA, and the LS (reviewed in Riters, 2012). Increased singing behavior is associated with increased neural induction in the VTA during both sexual and agonistic encounters (Bharati and Goodson, 2006; Heimovics and Riters, 2005; Maney and Ball, 2003). Lesion and electrophysiological studies link activity in the VTA to proper production of sexual song (Hara et al., 2007; Huang and Hessler, 2008; Yanagihara and Hessler, 2006), and LS lesions disrupt aggressive responses to territory intrusion (Ramirez et al., 1988).

As in birds, the context-independent association we observed between activation in VTA and LS homologues and behavior may reflect evaluation of social stimuli and motivation to respond to these stimuli generally. Parallel patterns of activity and association with behavior in TPp and Vl are consistent with the hypothesis that connections between these brain regions play a role in mediating their activity and, by extension, behavioral output (Maeda and Mogenson, 1981). Importantly, our observation that TPp and Vl activation are positively associated with behavioral activity regardless of context suggests that individual behavioral differences may stem largely from differences in motivation that are independent of social context. Thus, integration of social, other external, and internal cues, may be able to modulate TPp and Vl activity to modify behavioral motivation across social as well as other behavioral contexts.

### Conclusions

Although male guppies performed similar behaviors in competitive versus mating contexts, we observed distinct patterns of activation associated with social context and behavioral output at the neural level. Induction of pS6 in the POA differed based on the social context male guppies experienced, while pS6 activation in TPp and Vl was positively associated with behavioral output. Taken together, the patterns we observe support a model in which distinct aspects of behavior are mediated by a balance of activity across distributed nodes of the SDMN (Teles et al., 2015), such that activity in distinct brain regions reflects behavioral context versus social motivation in a species in which behavioral repertoires are shared across social contexts.

## Acknowledgements

We thank the members of the Guppy Lab for help with fish care.

## Data availability

The guppy brain atlas is available in the Supplementary Material associated with this manuscript and on our lab website. Raw data are available upon request.

---

**Figure S1:**
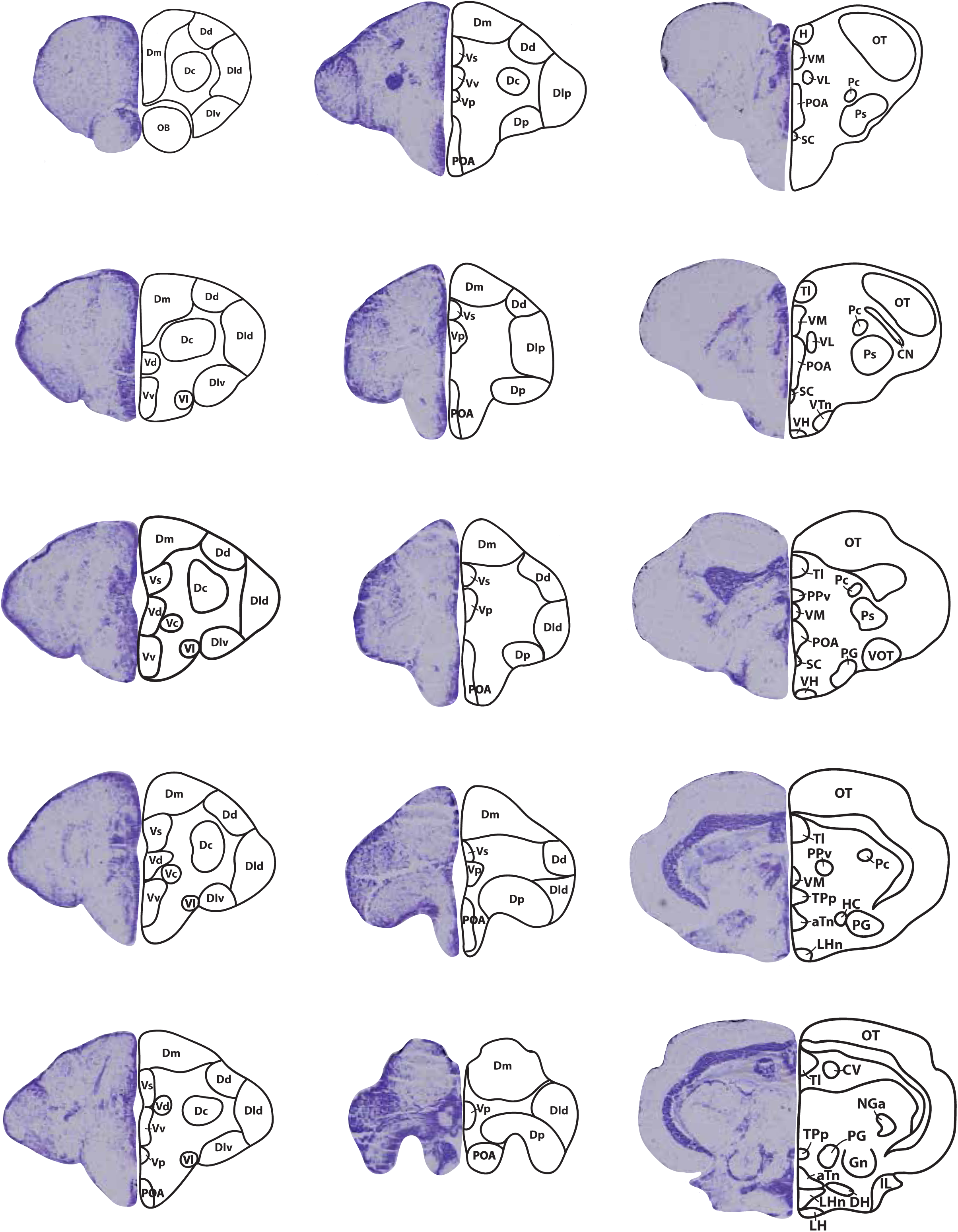
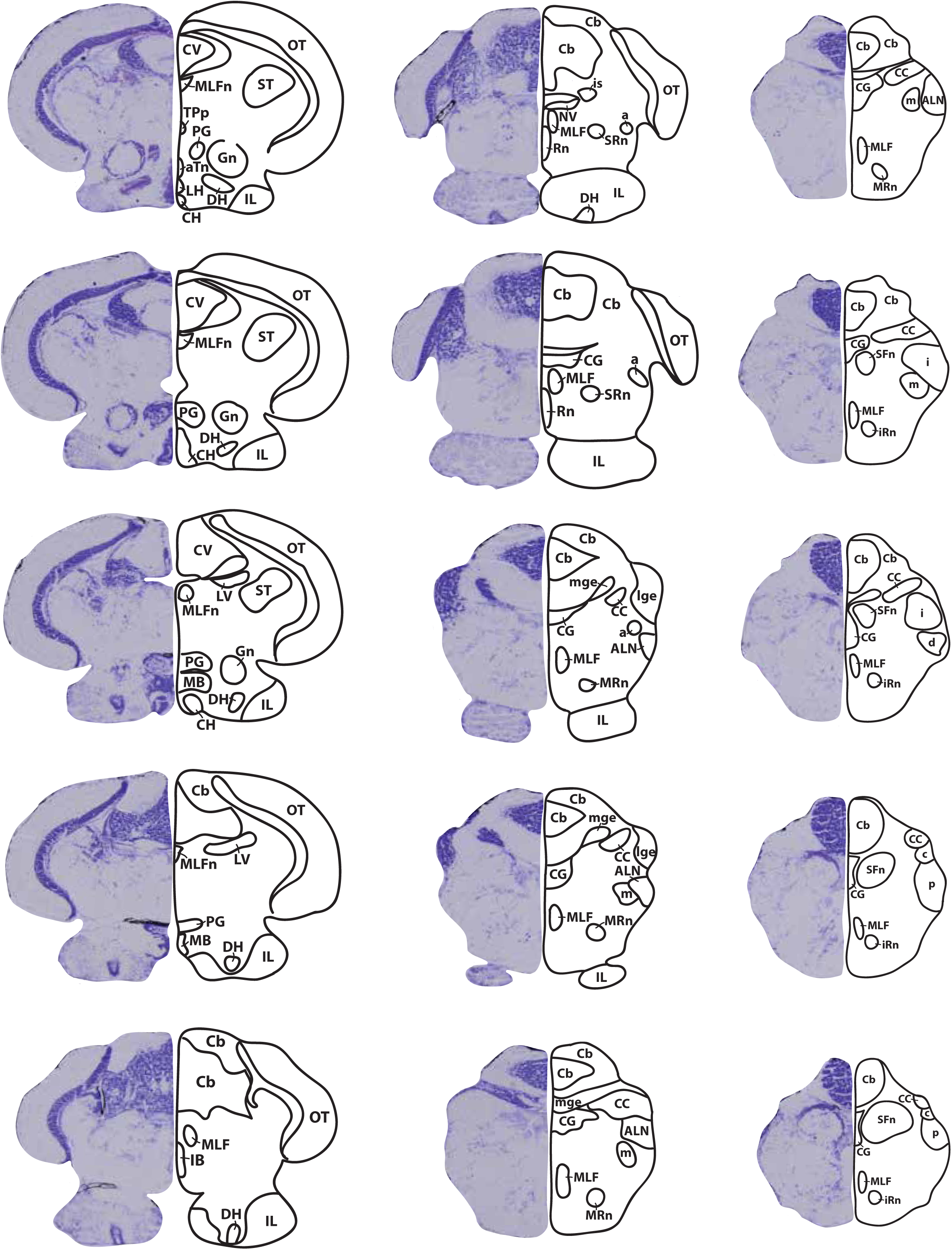
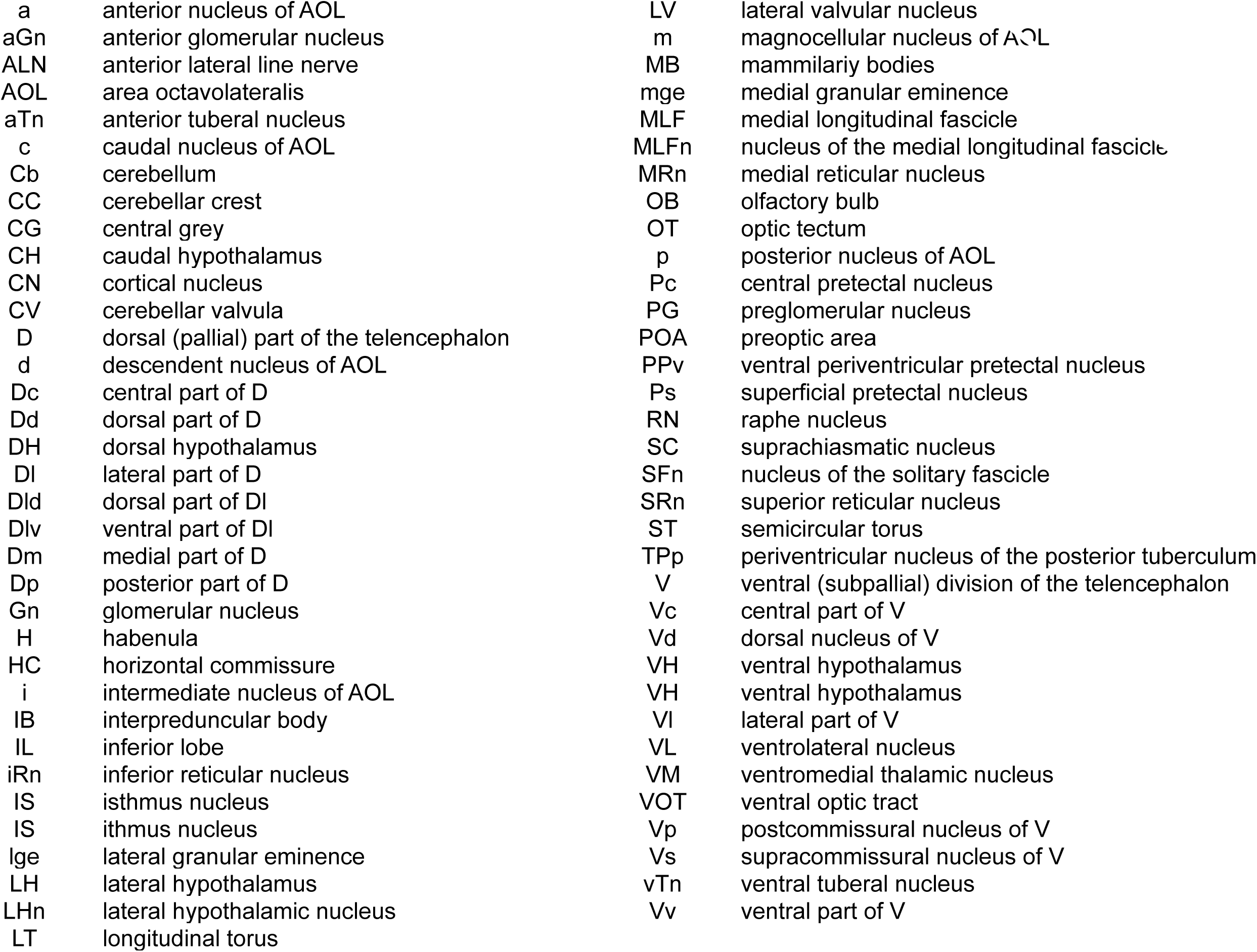
Neuroanatomical atlas of the guppy brain.

